# The Vector Competence and Vectorial Capacity of *Aedes albopictus* for Ross River Virus in the United States as a Function of Temperature

**DOI:** 10.64898/2026.01.09.698595

**Authors:** Joseph Spina, Donghyun Seo, Chang-Hyun Kim, Christopher Stone

**Affiliations:** Illinois Natural History Survey, Prairie Research Institute, University of Illinois Urbana-Champaign

**Author notes:** Joseph Spina 1816 Griffith Dr., Champaign IL 61820.

**Keywords:** Emerging viruses, vectorial capacity, vector competence, container-breeding mosquitoes, alphavirus, Culicidae

## Abstract

Ross River virus (RRV) is a medically significant arbovirus endemic to Australia where it causes an average of 5000 reported human cases per year. Due to concern for possible transmission outside of its endemic range, we examined the vector competence and vectorial capacity of RRV by *Aedes albopictus* (Skuse 1894) to contribute to risk assessments for outbreaks in the United States. We tested *Ae. albopictus* mosquitoes exposed to RRV at three temperatures and five time points post-infection for infection and dissemination. We also collected saliva from exposed individuals incubated at the same three temperatures to determine the presence of viral RNA. Additionally, we assessed the impact of viral infection on life expectancy as a function of temperature. Rates of infection were unaffected by time and temperature and 89.3% of individuals were infected. 85-100% of infected individuals displayed dissemination by day 7 at all temperatures. The extrinsic incubation period as determined by the saliva collection was approximately 6 days and 78.6-100% of individuals tested expressed salivary viral RNA at some point in the experiment which is comparable to the rates of overall infection. The vectorial capacity of RRV with *Ae. albopictu*s as a vector ranged from 10.23-23.37 indicating that these mosquitoes are competent potential vectors for RRV. Due to prior outbreaks RRV has caused outside of its endemic range, these findings warrant further investigation in the transmission potential of RRV in the United States.

**Author Summary:** The last 30 years has seen a steady increase in mosquito-borne illness globally, much of which has been driven by viruses such as dengue establishing in new areas as a consequence of the spread of its primary vectors and environmental changes. In order to mitigate the continued increase in the prevalence of mosquito-borne illness, it is necessary to determine which viruses are likely to spread beyond their native range. Ross River virus (RRV) is a mosquito-borne virus which, while nonlethal, commonly causes a litany of debilitating side-effects for extended periods of time. RRV is endemic to Australia, however the recent introduction of a known RRV vector to California as well as the widespread prevalence of a suspected RRV vector, *Ae. albopictus*, across the country, could prime RRV for an outbreak in the continental United States. As part of a risk assessment for the outbreak potential of RRV in the continental United States, we investigated the vector competence and vectorial capacity of *Ae. albopictus*. We find that this species is a competent vector for this virus across a range of relevant temperatures.

## Introduction

Ross River virus (RRV) is a single-strand positive-sense RNA virus in the Semliki Forest virus complex of the genus *Alphavirus* in the family Togaviridae. It is closely related to other medically significant arboviruses in the Semliki Forest virus complex such as Chikungunya virus, O’nyong-nyong virus, and Mayaro virus (6; 31; 26; 24; 1). RRV is endemic to Australia and is the causative agent of Ross River fever which is the most medically significant vector-borne disease in Australia with an average of 5000 annual clinical human cases reported (27). This represents a prevalence of 25.7 cases per 100,000 population based on the population of Australia at the time of reporting (2), higher even than the 20 cases per 100,000 population of Lyme disease in the United States (18).

Ross River fever is a nonfatal illness and clinically resembles other vector-borne diseases with headache, rash, and fever as common symptoms (5). However, Ross River fever is a particularly debilitating illness due to the occurrence of severe arthralgia (joint pain and swelling) in upwards of 83% of all cases, and which persists for more than six months in half of all instances (5). Due to the severity and long-lasting nature of these symptoms, the prevalence of the illness, and a lack of approved vaccines and antiviral therapies (20), RRV is highly significant from a public health and economic standpoint in Australia.

The transmission ecology of RRV is characterized by its considerable putative host and vector diversity. RRV has been detected in over 40 field-collected mosquito species across multiple genera and over a dozen mammal species have been implicated as potential reservoir hosts (27). The reservoir hosts of RRV were long-assumed to be marsupials such as kangaroos, wallabies, and possums. However, recent epidemiological, serological, and laboratory data has implicated a wider range of hosts including placental mammals including horses (*Equus)*, brown rats (*Rattus)*, flying foxes (*Pteropus*), and humans (27; 20). Furthermore, there have been major outbreaks in Australian cities including Perth and Brisbane, as well as on neighboring Pacific islands including Fiji and Samoa (27; 19). These have occurred in areas of high human population density despite a low abundance or absence of marsupial reservoir hosts, indicating that humans may be a more important host than has previously been recognized (13).

Humans express high viral titers for 1-6 days during an RRV infection (15). Furthermore, there are several container-breeding *Aedes* vector species in areas affected by RRV (namely Australia and Fiji) including *Aedes aegypti* and *Aedes albopictus* (27). These species coexist with humans in large numbers all over the world, and they are likely to become more widespread as the climate warms (15; 16). This raises the concern that additional areas outside the current endemic range may be vulnerable to RRV outbreaks. In fact, the outbreak in 1979 which led to over 500,000 cases in the Pacific is believed to have been initiated by a single viremic traveler to Fiji from Australia which lends credence to this idea (19).

The potential public health and economic tolls posed by RRV and developments in our understanding of RRV’s ecological dynamics with regards to its vectors and hosts warrant a risk assessment for the introduction of this virus in other parts of the globe (27). *Ae. albopictus* is a container-dwelling mosquito species which has been implicated as a potential vector in urban RRV outbreaks (3). This species has a large introduced range in the continental United States, generalist feeding habits with a high affinity for human bloodmeals in certain environments, and a reputation as a known vector of a number of other arboviruses, including alphaviruses such as chikungunya (12; 35). The ectothermic nature of mosquitoes means that atmospheric temperature plays a major role in their life and vector characteristics (25). An important consideration is therefore how the potential intensity of transmission in new regions depends on temperature, including seasonal temperature variation, climate change, or urban heat effects.

Vectorial capacity is a metric of the transmission potential of a given vector population for a given pathogen, which represents the mean number of secondary host infections resulting from a transmission cycle from one initial host (10). It also serves as a relative measure to determine when transmission of a given pathogen is likely to be most intense as well as what variables result in the strongest variations in transmission intensity. The vectorial capacity equation consists of biotic factors and life history traits, can incorporate vector competence, and can be formulated to integrate abiotic factors such as temperature (16). Important components include the probability of a mosquito surviving the extrinsic incubation period of the pathogen in question, the number of infected lifetime bites made by mosquitoes that have survived the incubation period, as well as the number of host-seeking mosquitoes per host per day. Due to the complex nature of vector-borne disease, this metric provides a useful and mechanistic lens through which to interpret a mosquito’s capacity for spreading a given virus in a particular environment. As such, we assessed the vector competence and life history traits of *Ae, albopictus* to determine the vectorial capacity for RRV as a function of temperature. The vector competence of RRV has already been assessed in Malaysian *Ae. albopictus* mosquitoes. However, the competence characteristics of populations of the same species can vary greatly by geography as well as temporality, with notable differences existing even between populations derived from the same region in different years (11)(14). Furthermore, the risk of an RRV outbreak in the United States has yet to be assessed, making an analysis such as this important for public health readiness for vector borne disease in the United States.

## Methods

The current study was conducted using 5^th^ generation adult *Ae. albopictus* originating from eggs collected in Champaign County, IL in 2023. We performed a vector competence analysis for RRV in order to determine rates of infection, dissemination, and salivary transmission, as well as a survival assay to determine whether RRV infection negatively impacts life expectancy in female mosquitoes.

### Mosquito rearing and propagation

*Ae. albopictus* eggs were acquired at two sites with five collection locations each at two time points in Champaign County in 2023. These eggs were passaged until fifth generation eggs were collected and desiccated until needed for experimentation.

### Viral culturing

Stocks of Ross River virus were strain T48 (GenBank accession number GQ433359) and were obtained from BEI resources (Manassas, VA). The virus was originally obtained from human serum in March 1980, and propagated in C6/36 *Ae. albopictus* larval epithelial cells by BEI. After acquiring the virus, it was passaged twice in Vero cells, frozen at -80 °C, and an aliquot was thawed and quantified via plaque assay. The frozen viral aliquots had a post-thaw concentration of 1.186E9 PFU/mL. This plaque assay protocol was adapted from a protocol for West Nile virus quantification (22).

### Mosquito rearing and infection

In preparation for infection, preserved F5 egg papers were placed into laminated steel 22.5×27.5cm pans with 1000 mL of hatching solution composed of DI water and 125 mg bovine heart and brain infusion. After 24 hours, hatched larvae were transferred to separate pans containing 1000 mL DI water at a concentration of 100 larvae per pan. 30 mg of finely-ground Tetramin brand fish food was added to each pan on Day 0 and no food was added on Day 1. 45 mg of Tetramin was added on Day 2 and 60 mg was added each day thereafter until all larvae had pupated. Upon pupation, the pupae were transferred to a cup of DI water in a mesh-lined 4 liter paper deli container and provided a 10% sucrose solution upon emergence. Each cage held a total of 150 adult male and 150 adult female mosquitoes. The F5 mosquitoes were left at 28 °C and 70-85% relative humidity for 5-10 days to allow all individuals to emerge from their pupal state and mate.

For an infectious blood meal, bovine blood with 2.02 × 10⁶ plaque-forming units (PFU) per mL of RRV was freshly prepared by mixing 10.1 μL of a 1.2 × 10^9^ PFU/ml RRV suspension with 6 ml bovine blood in citrate (Hemostat Laboratories, Dixon, CA). The RRV concentration in a blood meal is considered a plausible titer of a viremic human (15). Two milliliters of the prepared infectious blood meal were used to feed 5-10-day-old mosquitoes in a cage via a Hemotek blood-feeding system (Blackburn, United Kingdom), and mosquitoes were allowed to feed for one hour. Experiments requiring negative control mosquitoes had the same blood feeding protocol with the same colony conditions albeit with bovine blood in citrate which was not treated with viral supernatant.

After feeding blood-fed mosquitoes were sorted after cold-anesthetization, individually housed in 500 ml paper deli containers with plastic lids featuring a center hole covered with mesh. A total of 525 were blood-fed, and separated at random into three groups of 175 cages based on incubation temperature schemes. One group each was placed into the three incubators set to 20 ℃, 24.5 ℃, and 29 ℃ (Percival Scientific, Perry, IA) under a 16:8 light-dark cycle. Each cage was provided with a cotton ball soaked in a 10% sucrose solution every two days, and mosquito survivorship was monitored daily during the incubation period. The relative humidity in each environmental chamber was 80-90%. The selected temperatures represent the average high temperatures during early, middle and late mosquito seasons in Cook County, IL (33), an area at the northernmost boundary of *Ae. albopictus*’s current range in the US.

### The effects of temperature on RRV infection and dissemination rates

A subset of 25 cages with surviving exposed mosquitoes was selected randomly from each temperature scheme at different incubation times (2, 7, 10, 13 and 16 days post-infection (DPI)) and utilized for assessing RRV infection. Upon harvest at the designated DPI, the mosquitoes were killed by freezing at -80 °C and then dissected on dry ice to separate the head, legs and body, which were stored in separate tubes at -80 °C until nucleic acid isolation. The head and legs served as two independent measures of viral dissemination, while the bodies were used to determine infection rates.

### The assessment of mosquito saliva for RRV transmission

A subset of 45 fully engorged adult mosquitoes, fed with an RRV-infectious blood meal, was divided into three groups of 15 females. Each group was placed in a 50 mL conical tube containing a 2.3 cm circular filter paper soaked in a 50% sucrose solution and kept at one of three temperature settings (20 °C, 24.5 °C and 29 °C) for 16 days. The tubes were modified to allow easy replacement of the filter paper without losing mosquitoes. The filter paper was replaced every 24 hours, and each harvested filter paper was placed in a 2 mL microcentrifuge tube and stored at −80 °C until nucleic acid isolation. This protocol was adapted from a saliva collection protocol to detect Chikungunya virus (7).

### The survivorship of RRV infected mosquitoes

Two additional subsets of 150 bloodfed adult mosquitoes were acquired, fed with either RRV-infectious blood meal or control bovine blood. These subsets were further divided into three groups of 50 females, maintained under one of three temperature settings (20 °C, 24.5 °C and 29 °C), and monitored daily for mortality. The experiment continued until all mosquitoes had died. Dead mosquitoes were removed on the day of death with their date of death being recorded as DPI.

### Molecular detection of RRV

RNA isolation was performed on a KingFisher Apex platform (Thermo Fisher Scientific, Waltham, MA). Protocols recommended by the manufacturer of the MagMAX Viral/Pathogen Ultra Nucleic Acid Isolation Kit (Applied Biosystems, Waltham, MA) were followed, with slight modifications at the tissue lysis step. For mosquito tissue samples, tissues were lysed using a TissueLyser II (Qiagen, Germantown, MD) after being placed in a Type D bead tube (Takara Bio, San Jose, CA) containing a lysis buffer composed of 288 µL Buffer AVL (Qiagen, Germantown, MD) and 12 µL 1 M DL-Dithiothreitol (DTT; Sigma-Aldrich, St. Louis, MO).

For filter paper samples, the TissueLyser II was inefficient. Instead, RRV particles were separated from the filter paper by incubating it in 500 µL of lysis buffer composed of 480 µL Buffer AVL and 20 µL DTT for 30 minutes at 50 °C.

Two hundred microliters of the resulting supernatant were loaded onto the KingFisher Apex system and processed for RNA isolation using the MagMAX Viral/Pathogen Ultra Kit (Applied Biosystems, Waltham, MA). The final eluted RNA volume was 70 µL.

RRV detection and quantification were performed using one-step quantitative real-time reverse transcriptase PCR (RT-qPCR) with the probe and primer set developed by Taylor et al. (32), targeting the NSP-3 (nonstructural protein 3) coding region of the RRV genome. The forward primer sequence was 5′-CCG TGG CGG GTA TTA TCA AT-3′, the reverse primer sequence was 5′-AAC ACT CCC GTC GAC AAC AGA-3′, and the FAM-labeled Minor Groove Binder (MGB) probe sequence was 5′-ATT AAG AGT GTA GCC ATC C-3′. RT-qPCR was performed in a 20 µL reaction volume containing 1× QuantiTect Probe RT-PCR Master Mix (Qiagen, Germantown, MD), 0.2 µL QuantiTect RT Mix, 1 µL each of 10 µM forward and reverse primers, 0.5 µL of 10 µM probe solution, 5.3 µL dH₂O, and 2 µL RNA sample. Thermal cycling conditions on the QuantStudio 5 (Applied Biosystems, Waltham, MA) were: 1 cycle of reverse transcription at 50 °C for 20 minutes, 1 cycle of DNA polymerase activation at 95 °C for 10 minutes, followed by 40 cycles of denaturation at 95 °C for 30 seconds and annealing/amplification at 60 °C for 60 seconds. Results were obtained as Cq values after setting a threshold line at 0.2, and a Cq value of ≤37 was considered positive. For quantification, a standard curve was constructed using RT-qPCR under identical conditions with a plasmid containing the PCR primer-flanking region of RRV, for which copy number was calculated.

### Statistical analysis

#### The effects of temperature on RRV infection and dissemination rates

A generalized linear model was used to determine whether temperature, time, or a combination of these factors influenced rates of infection and dissemination. A quadratic regression was used to model the growth and reduction of mean viral titers in the body samples throughout the course of the experiment. A logistic curve was used to determine the growth of viral titers in the dissemination samples derived from positive corresponding body samples. A Wilcoxon signed rank test was used to determine if there was a difference between the viral titers of the paired dissemination samples, and individuals with only one tissue sample testing positive were excluded from analysis.

#### The assessment of mosquito saliva for RRV transmission

The rate of salivary transmission was reported as a proportion of mosquitoes which had expressed viral RNA through their saliva for at least one time point. The time at which 50% of all mosquitoes in a given incubator had expressed RNA through their saliva was used to determine a probable extrinsic incubation period. Individuals which died within five days post infection were excluded from the final analysis.

#### The survivorship of RRV infected mosquitoes

Cox Proportional Hazards models were used to determine whether there were significant differences between the life expectancies of individuals in the experimental and control groups at the different temperature levels. The same test was used to compare the three experimental groups to one another to determine the effect of temperature on survival.

#### Vectorial capacity

Overall rates of infection and salivary transmission, extrinsic incubation period, and hazard ratios (HRs) obtained from the survival assay were combined with data from the literature to determine the vectorial capacity for a given temperature. The vectorial capacity model used is adapted from one originally published by Garrett-Jones and Shidrawi (10).

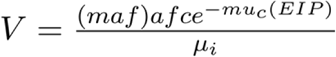

Where *m* refers to the average density of mosquitoes per human, *a* the daily rate of blood feeding, and *f* the proportion of bites taken on humans (and *maf* represents the number of female mosquitoes taking a blood meal per person per day, which can be informed by both human landing catch data and the proportion of bloodmeals taken from a human host), *c* represents the proportion of mosquitoes which will transmit the infection assuming they have consumed an infectious bloodmeal, *µ* is the daily mortality probability of a female mosquito with the subscripts *c* and *i* referring to uninfective/exposed and infective females, respectively, and EIP is the extrinsic incubation period. An overview of the variables, as well as the parameter values used, is provided (Table 1). This equation operates on the simplifying assumption that humans are the only potential RRV hosts and *Ae. albopictus* the only potential RRV vectors present in the continental United States.

The daily rate of bloodfeeding was estimated based on the gonotrophic cycle time of *Ae. albopictus* as a function of temperature (4). Landing rate data was obtained from a researcher at the Illinois Natural History Survey in Champaign, IL based on data obtained in Champaign County, IL for *Ae. albopictus* landing on human hosts (Mackay et al, in prep.). The proportion of bloodmeals taken from human hosts was based on a review of *Ae. albopictus* feeding behavior in North America (9).

Data collected by Nur Aida et al. (23) which determined life expectancy under uncontrolled laboratory conditions for *Ae. albopictus* mosquitoes kept at a mean temperature of 29 °C served as the baseline measurement for life expectancy equivalent in the 29 °C control group in the survival experiment. This estimate was divided by the HRs for the infected individuals to obtain probable life expectancies for mosquitoes at each temperature, and the inverse of these life expectancies was taken to obtain the daily expected mortality rates for the vectorial capacity model.

Vector competence across three temperatures was estimated in this study as well as extrinsic incubation period at 20 and 24.5 °C. Due to a mass mortality event of the 29 °C individuals used in saliva collections, the value for extrinsic incubation period at 29 °C was estimated based on a study of Malaysian *Ae. albopictus* mosquitoes infected with RRV.

## Results

### Infection and Dissemination

*Ae. albopictus* females challenged with RRV-infected blood meals showed high rates of infection and dissemination. The 20 ℃ amplification rates ranged from 0.84-0.96 (mean ± SD = 0.904 ± 0.061), the 24.5 ℃ amplification rates ranged from 0.88-1.0 (mean ± SD = 0.888 ± 0.071), and the 29 ℃ amplification rates ranged from 0.84-0.96 (mean ± SD = 0.888 ± 0.044) as illustrated in Figure 1. These outcomes were not significantly affected by day (Binomial GLM, Z=-1.10, p=.270) or temperature [Z(24.5)=1.46, p(24.5)=0.143, Z(29)=0.078, p(29)=0.816], indicating that variability in the probability of infection was dependent on other factors that were not measured here.

**Figure 1:**
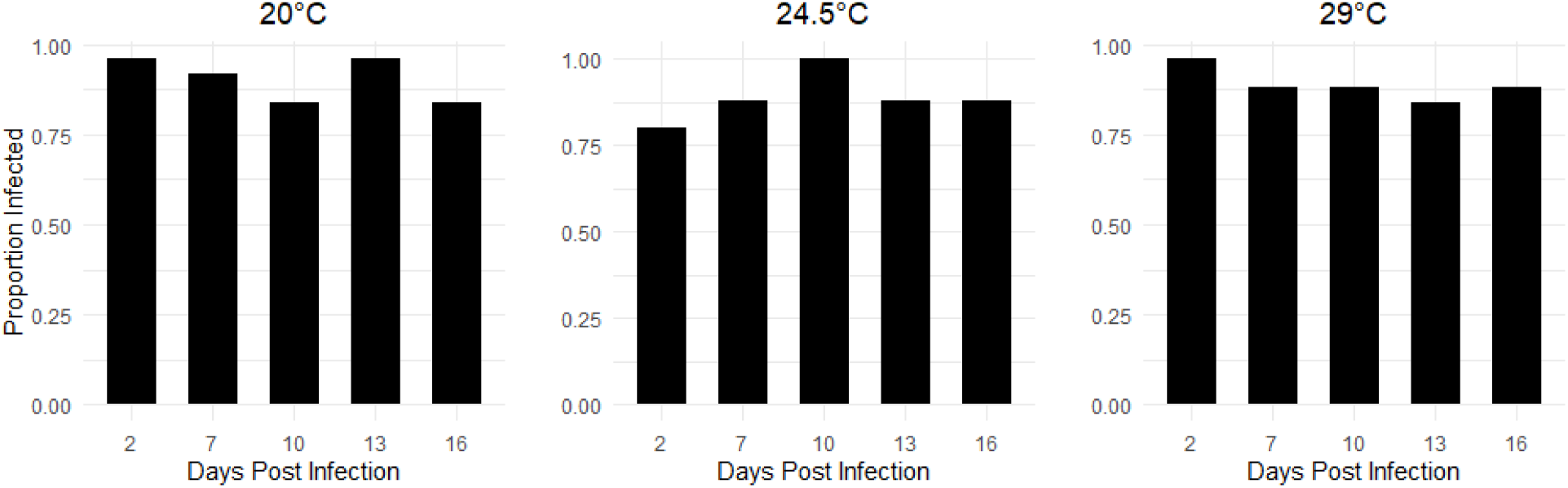
Rates of amplification of RRV by day post infection (DPI) and temperature. 20 ℃ mean=0.904±.061, 24.5 ℃ mean=0.888±.071, 29 ℃ mean=0.888±.0.44. Neither temperature nor time nor a combination of the two significantly altered infection rates. The overall mean infection rates across all groups was 89.3%.

The variation in mean titers of the body samples was described by quadratic regression models [R^2^(20)=0.276, R^2^(24.5)=0.378, R^2^(29)=0.275] as shown in Figure 2. The first and second order polynomial components were highly significant across all models (p(poly 1) < 0.001, p(poly 2) = 0.00129) indicating a strong linear component as a function of time and significant curvature in the data respectively.

**Figure 2:**
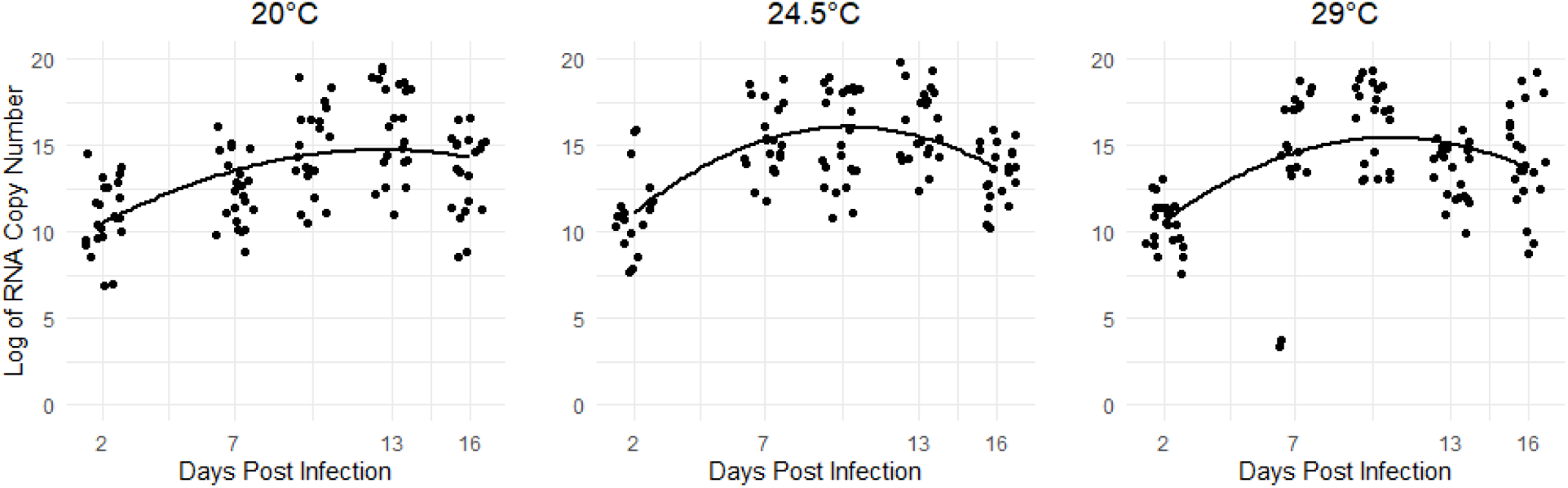
Changes in log-transformed viral titers in body samples by day and temperature modeled by quadratic regressions. RSE(20)=2.49, RSE(24.5)=2.20, RSE(29)=2.81. A gradual increase and decrease in mean viral titers over time can be observed at all temperatures.

The effect of temperature was significant in the 24.5 ℃ incubator relative to the 20 ℃ (T=2.39, p=0.017), but this was not the case in the 29 ℃ (T=1.10, p=0.280). When comparing the models to one another, the first order polynomial component was not significant in the 24.5

℃ group or 29 ℃ group relative to the 20 ℃ (p(24.5)=0.241, p(29)=0.477). The second order polynomial component was only significant in the 24.5 ℃ model relative to the 20 ℃ (p(24.5)=0.030, p(29)=0.102) indicating that there is only a difference in the curvature component of the 24.5 ℃ model relative to the baseline. This indicates that the rise and fall of the titers as well as their mean values did not differ significantly from one another except for the effect of temperature in the 24.5 ℃ model relative to 20 ℃.

There was no significant difference between the viral RNA titers of the paired head and leg samples from each individual (Wilcoxon signed rank test, W=186, p=0.118). As such, the leg samples were used for all dissemination analysis. Dissemination occurred gradually with less than 25% of individuals undergoing dissemination in each group by day 2, and dissemination rates of 75%-100% were the norm for all groups from day 7 onwards as shown in Figure 3.

**Figure 3:**
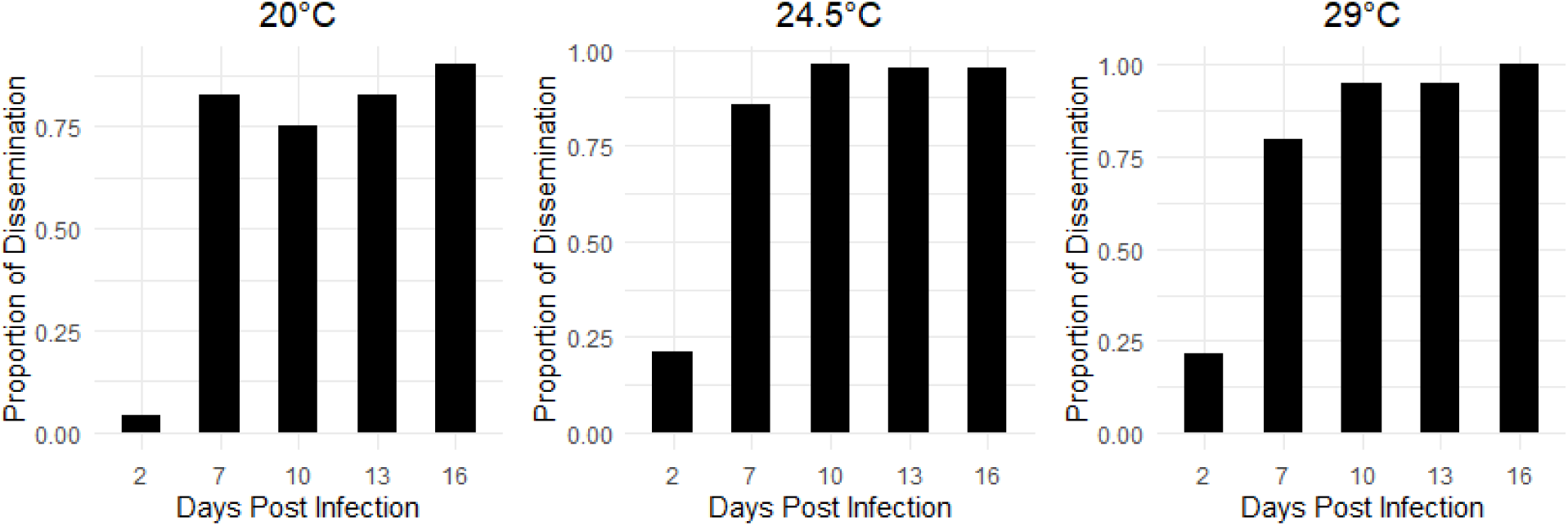
Rates of dissemination by day and temperature. Rates of dissemination were low at day 2 at all temperatures before approaching or reaching 100% by day 7.

A generalized linear model found no difference in rates of dissemination as a function of temperature in the 24.5 °C incubator (p =0.760) or the 29 °C incubator (0.845) when compared to the 20 °C. Time, however, was a reliable predictor of dissemination rates (p < 0.001). The logistic regression models fit the growth curves well for the 20 °C (McFadden R^2^ =0.444), 24.5 °C (McFadden R^2^ =0.377), and the 29 °C group (McFadden R^2^ (29)=0.595) as shown in Figure 4.

**Figure 4:**
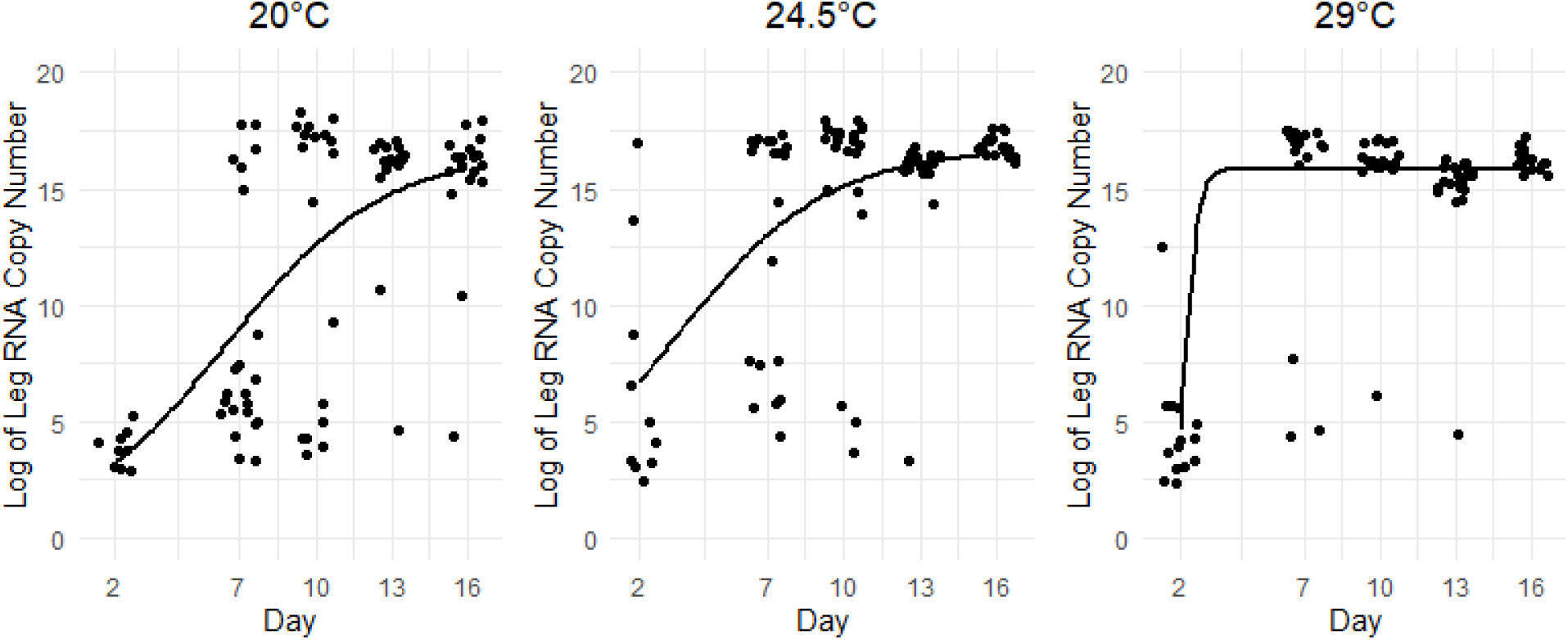
Changes in log-transformed viral titers in dissemination samples modeled by logistic curves. McFadden R^2^ (20)=0.444, McFadden R^2^ (24.5)=0.377, McFadden R^2^ (29)=0.595.

There were two apparent levels of RNA titers in the dissemination samples, one low level of approximately log(2.5)-log(7.5) and one high level of approximately log(15)-log(18) as shown in Figure 5. The log-transformed titer values excluding negative samples from day 2 to day 16 were compared using a Gaussian GLM. At 20 °C, mean titers rose from 1.51 [CI (0.124, 2.89)] to 14.82 [CI (13.3, 16.3)] (*p* < 0.001), at 24.5°C, mean titers rose from 3.64 [CI (2.12, 5.16)] to 16.9 [CI (15.4, 18.3)] (*p* < 0.001), and at 29°C, mean titers rose from 3.40 [CI (2.01, 4.78)] to 16.3 [CI (14.8, 17.7)] (*p* < 0.001). This elevation of dissemination titer values appears to have happened faster at higher temperatures with virtually all 29 °C samples exhibiting the higher titer value by day 7.

**Figure 5:**
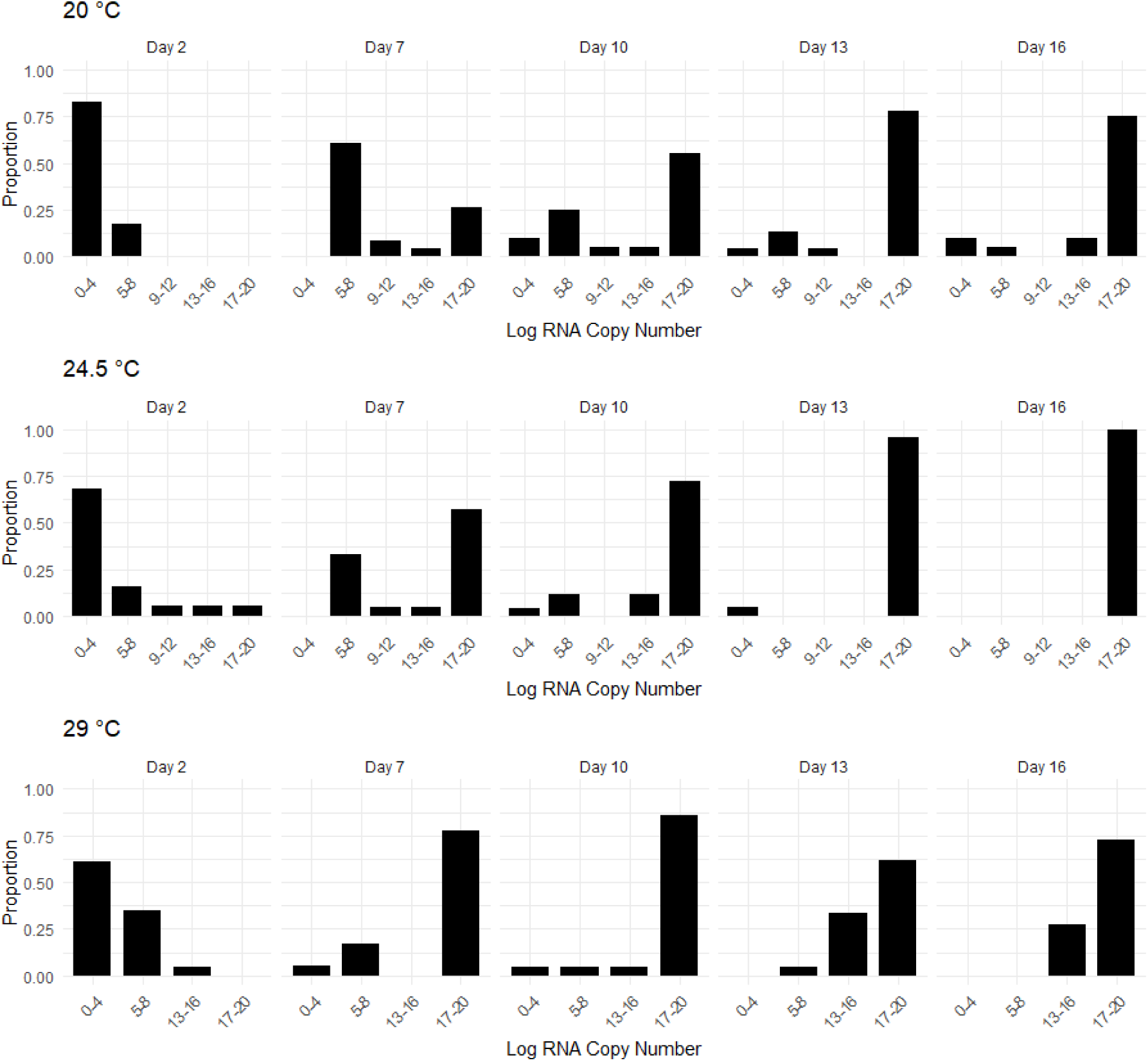
Distribution of log-transformed titer values in dissemination tissues by day and temperature. In any given group, a vast majority of samples had viral titers either in the log0-log4 range or the log17-log20 range.

### Salivary viral RNA transmission

The 29 °C mosquitoes experienced a mass die-off with a loss of 11/15 mosquitoes by day 5 that were found dead in the water cup. This group was excluded from the final analysis because we were not sure the cause of death. The earliest detection of Ross River virus in the 24.5 °C occurred three days post-infection while in the 20 °C group the first detection occurred five days post-infection. However, detection was uncommon in both groups until six days post-infection (Fig 6). We used the point at which at least 50% of individuals had expressed viral RNA through their saliva as an estimate of the extrinsic incubation period (28). Using this metric, the extrinsic incubation periods of both groups were 6 days, indicating that temperature did not significantly impact this value at 20 °C or 24.5 °C.

**Figure 6:**
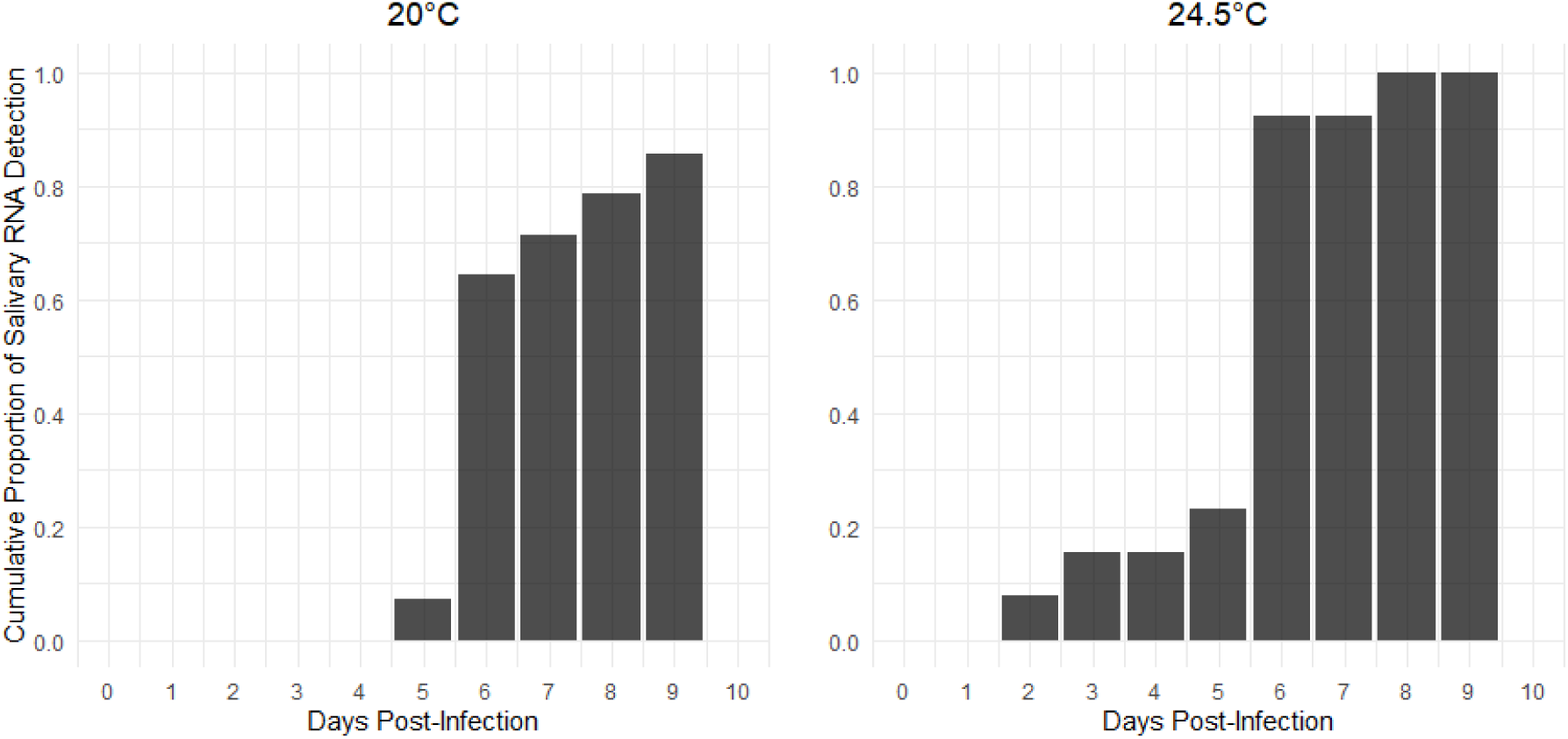
Cumulative proportion of mosquitoes which had expressed virus in their saliva by time and temperature. Early detections occurred in both groups, but the point at which 50% of individuals expressed virus was day 6 in both groups.

Eleven of the fourteen mosquitoes in the 20 °C incubator which survived at least five days expressed viral RNA in their saliva at some point during the experiment for an overall expression rate of 78.6%. All thirteen mosquitoes in the 24.5 °C incubator which survived at least five days expressed RRV-positive saliva at some point during the experiment for an overall expression rate of 100%. These values are in line with the rates of overall infection as established in the infection and dissemination experiment, indicating that a mosquito which is successfully infected with RRV is highly likely to express virus in its saliva.

### Survival

When contrasting survivorship between infected mosquitoes and uninfected controls at different temperatures, the Cox proportional hazards (PH) models determined that RRV infection had no significant impact on the mortality rate of mosquitoes kept at 24.5 °C [HR = 0.793, CI = 0.527-1.19, p = 0.264] or at 29 °C [HR = 0.819, CI = 0.548-1.23, p = 0.331]. There was, however, an increase in mortality rates of the infected mosquitoes compared to the control mosquitoes in the 20 ℃ incubator [HR = 1.66, CI = 1.07-2.56, p = 0.023] (Fig 7).

**Figure 7:**
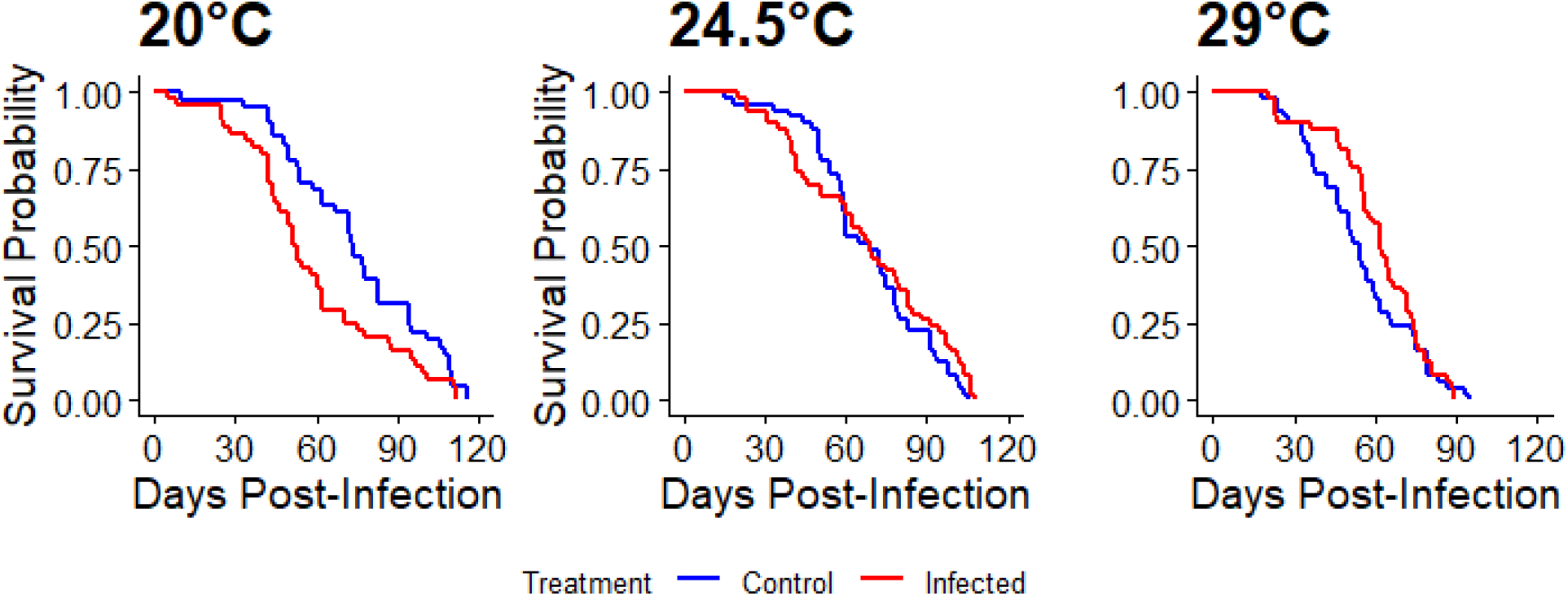
Cox proportional hazard models for daily survival probability in exposed versus control mosquitoes. There was a significant increase in mortality probability in exposed individuals at 20 °C, although this effect was absent at higher temperatures. [HR(20)=1.66, CI(20)=1.07-2.56, p(20)=0.023, HR(24.5)=0.793, CI(24.5)=0.527-1.19, p(24.5)=0.264, HR(29)=0.819, CI(29)=0.548-1.26, p(29)=0.331]

To calculate daily expected mortality rates for the vectorial capacity equation, the three infected groups were compared to the 29 ℃ control mosquitoes, which resulted in hazard ratios of 0.785, 0.483, and 0.635 for the 29 ℃ infected, 24.5 ℃ infected, and 20 ℃ infected mosquitoes respectively.

### Vectorial Capacity

Based on the vectorial capacity model (equation 1), we found estimates of RRV transmission potential in *Ae. albopictus* of 10.2, 23.4, and 14.7 at 20 ℃, 24.5 ℃, and 29 ℃ respectively. This indicates that Illinois-derived *Ae. albopictus* mosquitoes have the potential to cause onward transmission in case of an RRV introduction across a range of seasonally- and geographically-relevant temperature ranges, and that intermediate temperatures favor higher transmission rates.

## Discussion

Arboviruses have complex ecologies compared to most other viruses and need to be adapted to both hosts and vectors. As such, determining the vector competence of local mosquito populations is a critical prerequisite to understanding the epidemic potential of a given virus in a given area. Furthermore, due to the ectothermic nature of vectors, both entomological and environmental factors affect transmission intensity in a given area. Vectorial capacity is one metric which can factor in both entomological and environmental variables, and as such it is well-suited for determining the potential transmission intensity of outbreaks (17).

There have been large, confirmed RRV outbreaks in Pacific islands including an outbreak in 1979-1980 which affected over half a million people across American Samoa, Fiji, the Cook Islands, and Caledonia. These islands are not home to marsupials, and the outbreak is suspected to have been started by a viremic traveler (19), raising concerns about the potential for RRV to expand its range where potential hosts and vectors are present. Humans and horses have already been implicated as potential reservoir hosts and *Ae. albopictus* and *Ae. aegypti* have been implicated as potential vectors in urban outbreaks (27). Here we showed that *Ae. Albopictus* mosquitoes reared from eggs derived from central Illinois are highly susceptible to RRV infection, and a large proportion of infected females went to develop a disseminated infection as well as demonstrate salivary viral RNA expression.

It appears that RRV readily breaches the midgut of infected *Ae. albopictus* females and disseminates into the hemocoel after feeding on a bloodmeal with viral titers consistent with a human infected with RRV, and this occurs at high rates within 7 days post-infection. Titers in the leg and head tissues were distinctly bimodal, exhibiting either low or high levels of dissemination with few exceptions. Most individuals eventually reached the high titer-level of dissemination and these levels remained consistently high until the end of the experiment, although the timing at which we observed an increase in titers was temperature-dependent. Virtually all of the mosquitoes in the 29 ℃ incubators exhibited this elevated value by day 7 while this did not occur until days 10 and 13 in the 24.5 ℃ and 20 ℃ incubators respectively. This pattern was distinct from the estimated extrinsic incubation period of 6 days at both 20 ℃ and 24.5 ℃, where we did not see an influence of temperature. A noticeably shorter, three-day extrinsic incubation period was observed in Malaysian *Ae. albopictus* mosquitoes kept at 27-29°C and 70-90% relative humidity with a 12:12 light-dark cycle (8). The elevated dissemination titer values are similar to the titers observed in the body samples which could possibly be due to a gradual dissemination of virus where it takes several days after initial dissemination for viral hemocoel titers to reach equilibrium with the body samples. This bimodal pattern in virus titers was also documented in Malaysian *Ae. albopictus* and *Ae. aegypti* experimentally infected with RRV (8). This was suspected to be due to genetic diversity within mosquito or viral subpopulations. In the current experiment the virus had a single origin. The mosquitoes went through four generations in the lab, and although we did not measure genetic backgrounds present in this population, previous work on *Ae. albopictus* in central Illinois has shown the presence of multiple haplotypes (30). Once infected, 13 of 14 surviving individuals tested for salivary RNA in the 20 °C incubator and all 13 surviving individuals in the 24.5 °C incubator went on to express viral RNA through their saliva. This suggests a high level of competence for RRV in these mosquitoes. The rates of salivary transmission are comparable to overall rates of infection, pointing to a high level of transmission efficiency.

The survival assay supports the notion that temperature by itself is a determinant of longevity (4; 21). Surprisingly, we found that infection status affected mosquito longevity at low temperatures. Namely, at the lower temperature of 20 ℃ mosquitoes exhibited a significant fitness cost whereby the infected mosquitoes had a shorter lifespan compared to the control mosquitoes. At the higher temperatures, this effect was not evident. We did not observe a difference in mean body or leg viral titers among the temperatures, although there was a slower increase in viral titers at lower temperature. Future work on the immune response of *Ae. albopictus* by temperature levels, and whether these may exhibit a trade-off with longevity, would be of interest.

The vectorial capacity values for RRV in *Ae. albopictus* were 10.23, 23.4, and 14.72 at 20 ℃, 24.5 ℃, and 29 ℃ respectively. This suggests that *Ae. albopictus* is capable of causing onward transmission of RRV including in areas along the northern expansion edge in the continental U.S., such as Illinois. The model not only suggests that the vectorial capacity is highly variable based on temperature, but also that intermediate temperatures favor transmission, which roughly matches findings from a mechanistic modeling study which found peak transmission of RRV at 26.4 ℃ (29).

Our study has a number of caveats. We utilized a modified version of the saliva collection method developed by Fourniol et al. (7) which used honey-soaked filter paper. This method allows for the daily collection of saliva from individual mosquitoes over the course of the entire experiment. However, there is a certain amount of imprecision involved in this method. There is no way to guarantee that any given mosquito will feed on any given day. Furthermore, due to modifications made in the RNA extraction step, there was a high degree of RNA dilution relative to the RNA extraction protocol used for tissue samples which may have made the detection method less sensitive.

The vectorial capacity model primarily focused on temperature as a driver of variations in vectorial capacity. However, models such as this are particularly sensitive to variations in a vector’s host preference which itself depends on population density (34). As such, future analysis should consider such variations in order to build a more complete picture of RRV transmission potential in the United States. Future analysis should evaluate the vectorial capacity of *Ae. aegypti, Ae. notoscriptus*, and other possible vector species, as well as adjusting vectorial capacity as a function of human and mosquito population densities.

## Conclusion

Illinois-derived *Ae. albopictus* mosquitoes exhibit a high vector competence for RRV and vectorial capacity values that suggest the possibility of onward transmission across a range of temperature conditions, with a peak intensity at intermediate temperatures. These findings are notable due to the increase in the emergence of vector-borne diseases since the turn of the millennium, and this presents RRV as an arbovirus with the potential for emergence well outside of its endemic range. Further research on the likelihood of an outbreak of RRV in the continental United States resulting from the introduction of an infectious human during the *Ae. albopictus* vector season is warranted.

## Acknowledgements

We thank Aidan Berg for help with the statistical analysis, Kylee Noel for her help refining the saliva collection protocol, Jiayue Yan for reviewing and editing the manuscript, and Andrew Mackay for providing unpublished data. The following reagent was obtained through BEI Resources, NIAID, NIH: Ross River Virus, T-48, NR-51457.

---

**Supplementary Table 1:** Vectorial capacity model variables and values.

| Parameter | Meaning | Value(s) | References | Notes |
| --- | --- | --- | --- | --- |
| a(T) | Per-mosquito biting rate (gonotrophic time) | a(20) = .12345<br>a(24.5) = .222<br>a(29) = .286 | (4) | Gonotrophic cycle values (G(T)) from paper, biting rate estimated at 1/G(T) |
| c(T) | Vector competence | .893 | This paper |  |
| $\mu(T)$ | Adult mosquito mortality rate | $\mu_i(20C) = .032$<br>$\mu_i(24.5) = .025$<br>$\mu_i(29) = .040$<br>$\mu_c(20) = .0175$<br>$\mu_c(24.5) = .028$<br>$\mu_c(29) = .051$ | (23)<br>This paper | Life expectancy figures under uncontrolled conditions at mean daily temp of 29C derived from Nur Aida et al. and hazard ratios from survival assay applied to estimate 24.5C and 20C life expectancy for infected mosquitoes. |
| EIP(T) | Extrinsic incubation period | PDR(20) = 6<br>PDR(24.5) = 6<br>PDR(29) = 3 | (8)<br>This paper | 29C value from Fu et al.<br>20C, 24.5C estimated from this paper |
| MA | Landing rate | 24.25 | (Mackay et al, in prep.) | Mean landing rates of samples was 24.25., temperature did not play a role |
| f | Proportion of bloodmeals taken from potential hosts (.351) | .351 | (9) |  |

## Author Contributions

Joseph Spina - Conceptualization, data curation, formal analysis, Investigation, methodology, project administration, supervision, visualization, writing - original draft preparation, writing - review and editing

Donghyun Seo - Data curation, Investigation, methodology

Chang-Hyun Kim - Methodology, Resources, Supervision

Christopher Stone - Conceptualization, methodology, supervision, funding acquisition, resources, supervision

